# Introduction to Genomic Analysis Workshop: A catalyst for engaging life-science researchers in high throughput analysis

**DOI:** 10.1101/478024

**Authors:** Phillip A. Richmond, Wyeth W. Wasserman

## Abstract

Researchers in the life sciences are increasingly faced with the task of obtaining compute resources and training to analyze large, high-throughput technology generated datasets. As demand for compute resources has grown, high performance computing (HPC) systems have been implemented by research organizations and international consortiums to support academic researchers. However, life science researchers lack effective time-of-need training resources for utilization of these systems. Current training options have drawbacks that inhibit the effective training of researchers without experience in computational analysis. We identified the need for flexible, centrally-organized, easily accessible, interactive, and compute resource specific training for academic HPC use. In our delivery of a modular workshop series, we provided foundational training to a group of researchers in a coordinated manner, allowing them to further pursue additional training and analysis on compute resources available to them. Efficacy measures indicate that the material was effectively delivered to a broad audience in a short time period, including both virtual and on-site students. The practical approach to catalyze academic HPC use is amenable to diverse systems worldwide.

## Introduction

The era of genomics and DNA sequencing is being rapidly incorporated into life science research fields, spanning fields as diverse as population-scale human genetics, model organism studies and patient-focused precision medicine. Researchers have harnessed the wealth of data produced to unveil previously unattainable insights into their research questions. Although researchers across different fields are beginning to utilize the power of genomics, two major roadblocks of accessible compute resources and lack of training prevent them from being able to effectively analyze their own data (1). An emerging solution to deliver high performance computing (HPC) to researchers worldwide comes through centralization of compute resources, generally in the form of grid or cluster compute servers. Examples include Compute Canada which delivers coordinated grid computing to Canadian academic institutions, the National Computational Infrastructure that coordinates the supercomputer system Raijin for Australian researchers, XCEDE which coordinates compute power for academic researchers in science and engineering across the United States, and numerous existing European academic HPC solutions such as Partnership for Advanced Computing in Europe (PRACE) (2–5). Although such academic HPC systems have been historically used by researchers in physics and chemistry—fields dominated by high throughput calculations and big data computations—the systems are being increasingly used by life scientists. Even as funding for academic HPC systems grows, a lack of facilitation to introduce life scientists to their use remains a roadblock for many potential users.

The increasingly digital and quantitative analyses required in life science research has been noted for decades, but the arrival of accessible and affordable DNA sequencing technologies has accelerated the demand for skills that are not yet commonly incorporated into training programs. Thus, beyond the roadblock of HPC access and use, life science researchers must also acquire training in genomic analysis. Recent global surveys on the training needs for life science researchers in bioinformatics analysis revealed that most training comes at time-of-need or point-of-need, imposed by the necessity to analyze acquired data, instead of being a core part of the curriculum or formal education (6). The bioinformatics community has reacted to the challenge of meeting this time-of-need training on numerous fronts with great efficacy. Current training options in bioinformatics include massive open online courses (MOOCs) (7), static online tutorials (8,9) and in-person workshops (10–12). Although online forums provide a variety of resources, current tutorials are not compute-resource specific and require compute environment tailoring for HPC systems which is prohibitive to researchers lacking strong computational skills. Furthermore, surveys regarding efficacy place a high value on the practical analysis components, which are often not delivered in the online capacity due to difficulty of coordination (6). In-person workshops allow for hands on applications, but require overhead cost and scheduling, which can be a limitation to many researchers. Also, many of these workshops are primarily tailored for analysis on a laptop or desktop, which differ from the environment of HPC platforms (13). Moreover, some of the most popular workshops have transitioned away from introductory content, instead focusing on more complex topics and applications. Such workshops often assume background knowledge and experience in Linux and HPC. This transition occurred in part due to an expansion in online curriculum for introductory content, as well as the need to refine hundreds of workshop applicants to a feasible number of local attendees (12,13). Lastly, most available workshops, both online and in-person, are rigid in structure, not catering to the interdisciplinary and diverse skill levels prevalent in the genomic era. In summary, researchers in the life sciences are faced with the need for acquiring introductory analysis skills in an environment that has the capacity for high throughput analysis, and no current training options are singular and effective at delivering this training in a time-sensitive manner.

In light of the challenges, we sought to create a new educational approach to catalyze the use of academic HPC systems by life scientists. We envisioned a flexible, centrally-organized, easily accessible, interactive, and compute resource specific training for the foundational skills of genomic analysis. To this end, we implemented and taught a modular workshop series titled: “*Introduction to Genomic Analysis„*. The material was taught in both an online (using the Vidyo system) and local (at the BC Children’s Hospital Research Institute) capacity for a total of seven two-hour interactive sessions (14). We focused the content on practical application, effective use of available compute resources, and data analysis exercises, while avoiding other aspects of genomics such as theory and experimental design. Since these aspects of genomics are fundamental, we encourage students to utilize external open-source materials deposited in curated bioinformatics education repositories (8,15). The modularity of our workshop structure allows participants to pick-and-choose the sessions to attend based on prior experience level, thereby catering to both those new to the command-line and those experienced in Linux with an interest in exposure to genomic data analysis. Moreover, a strong emphasis was placed upon student evaluation in our workshop delivery. To benchmark student progress, problem sets were used as module exit and entrance requisites and a culminating exam assessed the ability to effectively analyze next generation DNA sequencing data.

In total, 80 participants attended at least one of the seven modules, and 58 certificates of completion were awarded based on completion of the core modules. Our initial cohort of students was diverse, including a spectrum of prior genomic analysis experience, equal gender representation, and various educational levels which recapitulates the world-wide audience (7). Successful completion of the workshop showed similar results for in-person and virtual attendees and across levels of prior experience. Post-workshop surveys of the course efficacy provided deeper insight into potential improvements for future implementations of this material, and expansion into more detailed non-introductory topics of analysis.

In summary, we demonstrate the utility of our workshop format and information delivery methodology with the hopes that other trainers across the globe will improve and adapt its content to fit the needs of the life-science research community. The materials will persist in an open source state, and future implementations of the workshop will be delivered as we continue to fill the gap in genomic analysis training.

In accordance with expectations and guidelines for effective bioinformatics workshops, the materials are all open source, stored online, and video-recorded for future use (16). All materials are published under Creative Commons ShareAlike 3.0 Unported License (CC BY-SA 3.0) and can be accessed at: https://phillip-a-richmond.github.io/Introduction-to-Genomic-Analysis/.

## Implementation

### Content Delivery

The materials for this workshop were designed to emphasize accessibility, consistency, modularity, and reusability, designed around an environment with capacity for high throughput analysis. We also place an emphasis on student evaluation by implementing assigned problem sets and a final exam.

#### Accessibility

The primary challenge for delivery was the presence of both local and virtual attendees, which allowed us to expand our attendance beyond typical locally-constrained workshops. We reached both audiences simultaneously by delivering the content through an audio-visual casting of the primary teacher’s screen, which showed both lecture slides and an open terminal for executing commands. Local attendees followed the session on a projector and had tables for their laptops, while virtual attendees tuned in via internet broadcast of the screen capture using Vidyo software. Teaching assistants (TAs) monitored both local and virtual participants, the later using openstack Etherpad, and responded to questions and provided assistance during the follow-along lecture (a lecture format in which students repeat commands as they are performed by the instructor). The Etherpad environment provides an online text document updated in real-time that contains links to resources, an attendance section, and a question-and-answer section monitored by the TAs (17). Lectures were recorded and made available after the session for participants who couldn’t attend or desired to re-watch the presentation.

#### Consistency

It was a goal in developing the materials to provide a consistent structure and process for each workshop session. Every session started with 45 minutes of a follow-along lecture that integrated commands for the audience to execute alongside the primary teacher. Commands executed were documented as GitHub Gists, which also contained additional details regarding command usage (18). At the end of the lecture, students spent 1 hour working through a problem set in small (2-3 person) groups with assistance from a roaming TA (virtually via Etherpad or locally in-person). An important introductory concept for genomic analysis is maintaining a well-organized hierarchical filesystem structure. To enforce this practice, which includes centralization of reference genomes as well as separation of raw from processed data, each session followed an identical file structure (Supplemental Figure 1). Within this structure, individual student directories titled with unique identifiers allowed for both tracking of student participation, and simultaneous use of common files throughout the session.

#### Modularity

Modularity was a key component of this workshop as attendees had different prior training and experience. To enforce modularity and allow participants to skip material they had previously mastered, we designed each workshop session with a prerequisite assignment that assessed the comprehension of the preceding material. The problem set at the end of each session would serve as the prerequisite entry to the next. We found this to be necessary due to a mixing of content between basic Linux command-line usage and applied short-read sequencing analysis. A final exam was performed after 4 core modules, which served as a comprehensive evaluation of basic skills necessary for genomic analysis in the HPC environment. Completion of the exam was necessary for attendance of the final three modules, which delved into more advanced topics.

#### Reusability

All course materials are available through open source licensing under Creative Commons ShareAlike 3.0 Unported License (CC BY-SA 3.0), and a single github-based website links together the relevant resources. These resources include the lecture slides, problem sets, Github Gists, course exam, Etherpad links, and recordings of the lecture and problem-set sessions. Additionally, the workshop directory on the HPC environment remains for future individual use within the structured environment.

#### HPC Environment

Contrary to numerous laptop-based learning modules and teaching practices, we designed our material around analysis within a HPC environment. The motivation was two-fold: 1) by teaching in this environment we could combine an introduction to computing and Linux with an introduction to genomic analysis; and 2) we prepared researchers to be comfortable with data analysis on the platform upon which they would analyze data in the future. We utilized a national Canadian grid HPC system set up with a module system for controlling software dependencies, running Torque-Moab scheduling software for distributing jobs from the head node to the compute nodes. With slight modifications, the material can be adapted for use on most academic HPC systems. We provided temporary guest accounts to participants lacking an account. This allowed us to expose the attendees to the environment and actively engaged them to utilize the resources that are made available to them as academic researchers. Numerous participants followed up with the systems administrators to acquire full accounts during the workshop and after its conclusion.

### Workshop Materials

Standards in workshop design and implementation have highlighted the importance of labelling learning objectives and prerequisites explicitly for each session (16). An overview of the workshop materials is displayed in Table 1. Since the workshop is modular, with benchmarks of understanding and analysis capabilities before and after each session, there is little redundancy in the per-session learning objectives. With this workshop design, we build on commands and topics from previous sessions with increasing complexity. For example, in Session 3, students learn about mapping short read DNA sequencing data to the human genome, and then, in Session 4, they re-run the same mapping commands but with the addition of variant calling. For students who are unable to master the prior material, they are encouraged to follow along and revisit the material from previous sessions. The workshop materials are available online through a coordinated website available at https://phillip-a-richmond.github.io/Introduction-to-Genomic-Analysis/.

**Table 1).** Workshop Materials. Breakdown of the workshop materials, including the learning objectives, commands learned, and prerequisites for each session.

| Session # | Title | Learning Objectives | Commands Learned | Prerequisites |
| --- | --- | --- | --- | --- |
| Session 1 | Introduction To Linux I: Basic Command Line | Basics of computing with high performance computers<br>Filesystem hierarchy<br>Basic command line operations<br>File handling & permissions<br>Standard out<br>File Formats: GTF, Clinvar Variant Summary | cp, ls, mv, cut, clear, mkdir, rm, wget, grep, more, less, head, tail, cat, gunzip, gzip, chmod | Log-in to the cluster |
| Session 2 | Introduction to Linux II: Interacting with the Queue | Basics of computing with high performance computers<br>Editing files with linux file-editors<br>Shell Scripts<br>Interacting with the queue<br>File transfer | emacs, nano, qstat, qsub, showq, module avail, module list, module load, scp | Session 1 Problem Set |
| Session 3 | Short Read Mapping and Visualization | Next Generation Sequencing Primer<br>Map short read DNA sequences to the genome using BWA<br>Convert file formats using Samtools<br>Utilize scheduler for pipeline execution<br>Visualize short read data in IGV<br>File Formats: SAM, BAM, indexed BAM, Fastq, Fasta | BWA mem, samtools sort, samtools view, samtools index | Session 2 Problem Set |
| Session 4 | Variant Calling (Small variants) | Exome sequencing<br>Mapped read post-processing<br>Variant calling for small variants<br>Variant compression and indexing<br>Visualization of variants and short-read data together<br>File Formats: VCF, compressed VCF, indexed VCF | freebayes, bgzip, tabix | Session 3 Problem Set |
| Session 5 | Variant Interpretation with GEMINI | Brief introduction to MySQL queries<br>Variant annotation with VEP<br>File Formats: PED file, gemini.db | gemini load, gemini query, gemini de novo | Course Exam |
| Session 6 | RNA seq I: Analysis with the Tuxedo Suite | RNAseq overview<br>RNAseq read mapping and visualization<br>Transcript assembly<br>Transcript quantification | HISAT2, Stringtie | Course Exam |
| Session 7 | RNA seq II: Differential Expression Analysis in R | R data loading and visualization<br>Differential expression using DESeq2 | R, DESeq2 | Course Exam |

## Evaluation of Efficacy

### Participant Breakdown

The group of participants in the workshop series was diverse in prior training, sex, academic standing, and mode of attendance (Figure 1A-C, Figure 2A). The academic standing of our participants was similar in distribution to that reported in global surveys of researchers interested in bioinformatics related training (7). The primary audience was graduate students actively completing their respective degrees, followed by undergraduates and post docs. Two faculty members attended the workshop, demonstrating the broad distribution in academic standing. Participants were evenly split between local and virtual (Compute Canada Vidyo screencast) attendees. This even distribution allowed us to draw conclusions regarding the efficacy of the teaching model where both local and virtual audiences coexist. Additionally, we had even splits in both prior genomic analysis experience and familiarity with Linux and HPC systems. Lastly, we had an equal representation of both sexes, which is an important factor in bringing equality to the field of computational biology, a historically male-dominated field(19).

**Figure 1).**
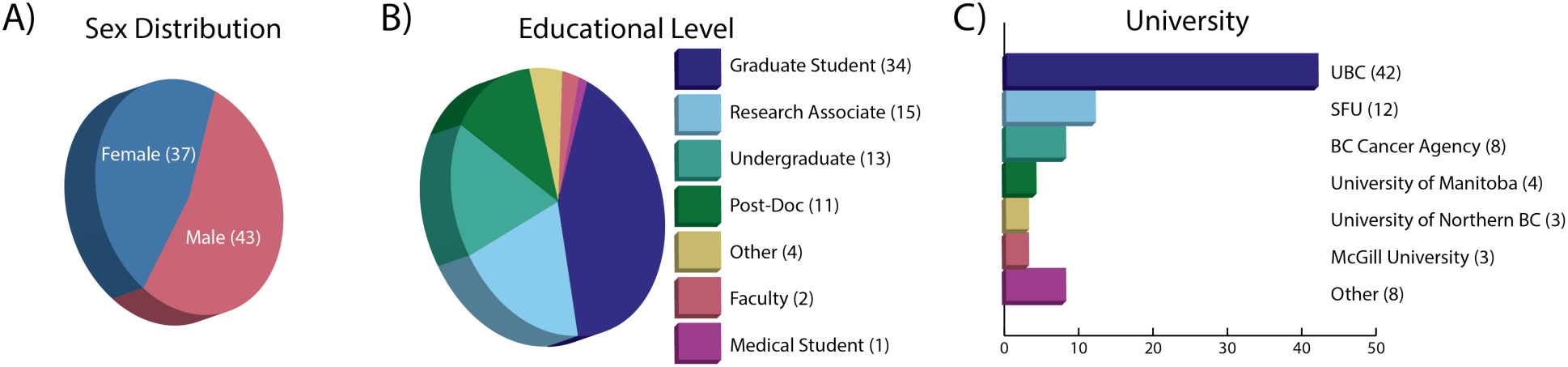
Participant Background. Description of workshop attendees including A) distribution of sexes; B) educational level; and C) the university from which they participated.

**Figure 2).**
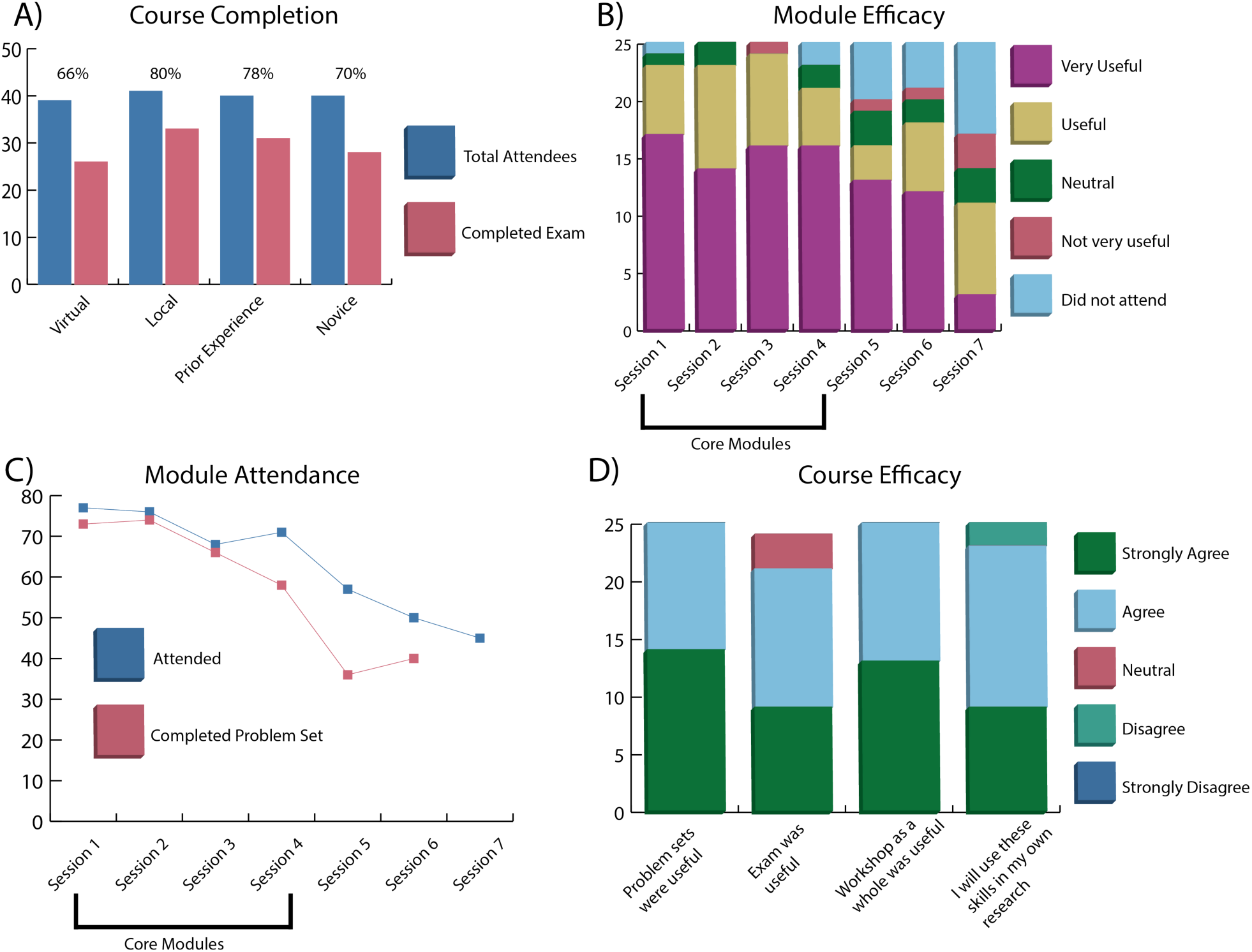
Workshop Results. A breakdown of the workshop results including A) distribution of course completion rates annotated by mode of attendance and prior experience; B) efficacy of each module based on survey responses; C) per-session attendance and problem set completion; and D) course efficacy breakdown including problem set and examination utility;

### Workshop Efficacy

Workshop efficacy was determined based on three measures: 1) the number of total participants versus the number of participants that completed the exam; 2) the per-session attendance and their ability to complete the problem set; and 3) a set of questions given in a post-workshop anonymous survey. Regarding course completion, 80 attendees participated in the workshop, 58 (72.5%) of which completed the exam. The completion rate was slightly higher for the local audience (80%) than the virtual audience (66%), possibly due to the stimulation of in-person collaboration between students which was noted as lacking in the post-workshop surveys from some virtual attendees (Figure 2A). When comparing between participants with prior and “zero” experience, the completion rates were surprisingly similar (Figure 2A), with both exceeding 70%. This represents a key success, as a key intention of the novel workshop design was engagement of researchers in the life sciences diverse levels of experience.

In analyzing the per-session attendance and problem set completion, we observed the efficacy of the module-based teaching methodology. While attendance was a measure of the interest in the subject matter, completion of the problem set identified student ability to effectively reuse the material taught during the first portion of the lesson on a unique data set. Sessions 1 through 3 had similar attendance and completion rates, and session 4 had a slightly lower completion rate since the problem set was also the mid-series exam (Figure 2 B,C). The exam covered material from session 1-4, and was more in depth and difficult than preceding problem sets. Attendance for the last three sessions was optional and based on participant interest which was reflected in the lower number of attendees. Problem sets for sessions 5 and 6 were not required for attendance of the following sessions, and therefore had a lower completion rate. There was no problem set for Session 7.

Lastly, we analyzed the survey responses from 25 attendees for both per-session evaluations and aggregated opinions about the utility of the content. Sessions 1 through 4 are the core modules that guided attendees through an introduction to Linux and the command-line, interacting with the scheduler and queue, basics of next generation sequencing (NGS) short-read mapping and visualization, and variant calling. These sessions scored well in both efficacy and attendance, and received favorable reviews (Figure 2B). Session 5 through 7, advertised prior to the workshop as “optional", went into more depth on subjects beyond the core NGS analysis including human genome variant interpretation and RNA-seq analysis. Despite an increase in the complexity of subject matter, the material was still judged to be accessible by the attendees. Some responses from the open commentary section were critical of the last few sessions, and suggested that those more complex topics need more time than allotted within a single 2-hour session. The progress assessments, including both problem sets and exam, were deemed useful by the majority of those that responded to the survey and received positive commentary throughout the workshop (Figure 2D). Lastly, 24/25 survey completers found the workshop useful and indicated an intent to utilize the materials and skills they learned in their own research projects.

## Future Perspectives

### Improvements

Constructive criticism of the workshop from attendees primarily focused on the final three modules, where advanced content was presented. In future iterations of the workshop, the initial 4 core modules will be taught as a set, while advanced topics will be offered separately. Those attendees that requested longer in-person sessions were referred to offerings from consortiums, such as Bioinformatics.ca, GOBLET, and ELIXIR, each of which provides excellent advanced in-person all-day workshops. Future iterations will test whether the advanced topics are viable to be taught in the practical skills focused format highlighted in this report.

On a more granular level, the problem sets were a key part of many student’s learning, but the system for delivering and grading those problem sets, via email and posting to locations on the shared server, was inadequate. Transitioning into a web-based platform (e.g. Moodle) for assignment delivery, completion, and grading, will allow for better feedback and communication regarding problem set questions.

### Following up with participants

#### If you don’t use it, you lose it

After being introduced to new concepts, a new language, and new compute environments, it is critical for continued practice to maintain and refine what you have learned. As a part of our workshop design, we can contact participants in the future to see how they progress in leveraging both the HPC resources and training received to process and analyze their own datasets. Our design of the workshop around the use of centralized compute resources allows us to engage with participants to see how effective the resources are for their research purposes. These long-term metrics will help inform retention and efficacy of materials, and identify gaps in training or resource needs that we can address through partnership with the academic HPC providers.

### Future workshops

We will deliver more workshops on both introductory and more advanced genomic analysis in the same modular format described above. By collecting similar efficacy metrics we can test how this workshop format performs with less introductory topics, and as part of a continued series. The overall goal is to establish training materials that can be delivered at time-of-need, that build strong foundations in genomic analysis utilizing the HPC systems. It is yet to be determined what the total attendee capacity is for this format, but our initial delivery reached 80 attendees, beyond the 30-40 person capacity of local workshops, and with the dual capacity of local and virtual delivery modes we anticipate audiences of over 100-200 participants.

## Conclusion

Training in bioinformatics and genomics will continue to be a critical component of the development of researchers in the biological sciences. Leaders in the research domain have proclaimed that acquiring computational analysis skills should be considered on par with learning the fundamentals of wet lab techniques (Sean Eddy blog post 2014). Currently, the lack of formal education in genomics and bioinformatics analysis for life-science results in numerous researchers seeking out time-of-need training to answer their research questions (6).

We have introduced a practical approach to the training of life scientists that focuses on getting researchers actively engaged with an academic HPC environment available to them for continued use beyond the confines of the workshop. The format incorporates both virtual and on-site participation, and the implementation successfully enables students with varying levels of experience to engage with the training at the relevant stages. The materials are available for re-use and are adaptable for use with most academic HPC architectures. As more centralized HPC systems become utilized and funded, we anticipate that this format of workshop will be invaluable in providing the foundation of training for researchers in the life sciences. Upon that foundation, researchers can further specialize their training needs and effectively participate in the high throughput technology revolution.

